# Common pitfalls during model specification in psychophysiological interaction analysis

**DOI:** 10.1101/2025.05.22.655642

**Authors:** Vicky He, Bahman Tahayori, David N. Vaughan, Heath R. Pardoe, Graeme D. Jackson, Chris Tailby, David F. Abbott, the Australian Epilepsy Project Investigators

## Abstract

Psychophysiological interaction (PPI) analysis is a widely used regression method in functional neuroimaging for capturing task-dependent changes in connectivity from a seed region. The present work identifies, and provides corrections for, common methodological pitfalls in PPI analysis that compromise model validity. Firstly, if the seed time series is extracted with prewhitening, the temporal structure of the signal is altered and subsequent deconvolution of prewhitened data becomes suboptimal. Furthermore, prewhitening again during model fitting results in double prewhitening of the seed regressor. Secondly, a failure to mean-centre the task regressor when calculating the interaction term can also lead to model misspecification and potentially spurious inferences. By using simulations and empirical language fMRI data from the Australian Epilepsy Project, we demonstrate the adverse effects of these issues, and how they are resolved when corrected. A systematic review of current practices revealed widespread model misspecification, and underreporting of methods, in published PPI studies. We provide clearer reporting guidelines, and advocate for appropriate methods for handling of prewhitening and mean-centring to ensure the validity of PPI analyses.

## 1 Introduction

Psychophysiological interaction (PPI) analysis is a multiple regression neuroimaging analysis method that estimates task-dependent changes in the functional connectivity from a seed region to other brain areas (Friston et al., 1997). Despite its widespread use, the correct specification of the PPI model has been a topic of considerable debate (e.g. Gitelman et al., 2003; O’Reilly et al., 2012; Di et al., 2017; Masharipov et al., 2024). In this paper, we focus on PPI analyses of functional magnetic resonance imaging (fMRI) data. We examine the implications of two model specification steps on model performance: (1) prewhitening and (2) mean-centring. We assess how these steps influence group-level analyses, including group mean comparisons and behavioural regressions.

In a system modelled by PPI, the relevant interactions of interest occur at the neuronal level rather than the level of the haemodynamic responses. To capture the neuronal response, and model its interaction with the psychological variable, one can deconvolve the haemodynamic response function (HRF) from the seed time course (Gitelman, 2003). This has been shown to increase sensitivity in PPI studies, especially in event-related designs (Masharipov et al., 2024; Di & Biswal, 2017).

The above approach can be carried out in the SPM software (Penny et al., 2006) and the generalised PPI toolbox (gPPI; McLaren et al., 2012). One typically first extracts a time series of a seed region using SPM’s time series extraction function. This extracts the first eigenvariate of the voxels in the region of interest by fitting a general linear model (GLM), so as to remove the effects of confound regressors such as realignment parameters. In this process, the extracted time series is prewhitened (see Fig. 1a) to ensure that when regressing out confounds, the errors are independent. Whilst prewhitening may be sensible to ensure valid inference at the confound correction stage, we argue that it adversely affects two downstream processes. Firstly, the subsequent deconvolution step will be applied to a prewhitened time series, potentially biasing the estimation of the underlying neuronal response. Secondly, the seed regressor that enters the PPI model design matrix is already prewhitened. Thus, when prewhitening is applied during the actual fitting of the PPI model, the seed regressor is effectively prewhitened a second time, causing it to depart from the seed time course it intended to capture (Fig. 1a). We refer to this as the double prewhitening issue. As a solution, we propose multiplying the extracted seed time series by the inverse of the whitening matrix (**W**^-1^) before downstream processing. We refer to this process as whitening inversion.

**Figure 1.**
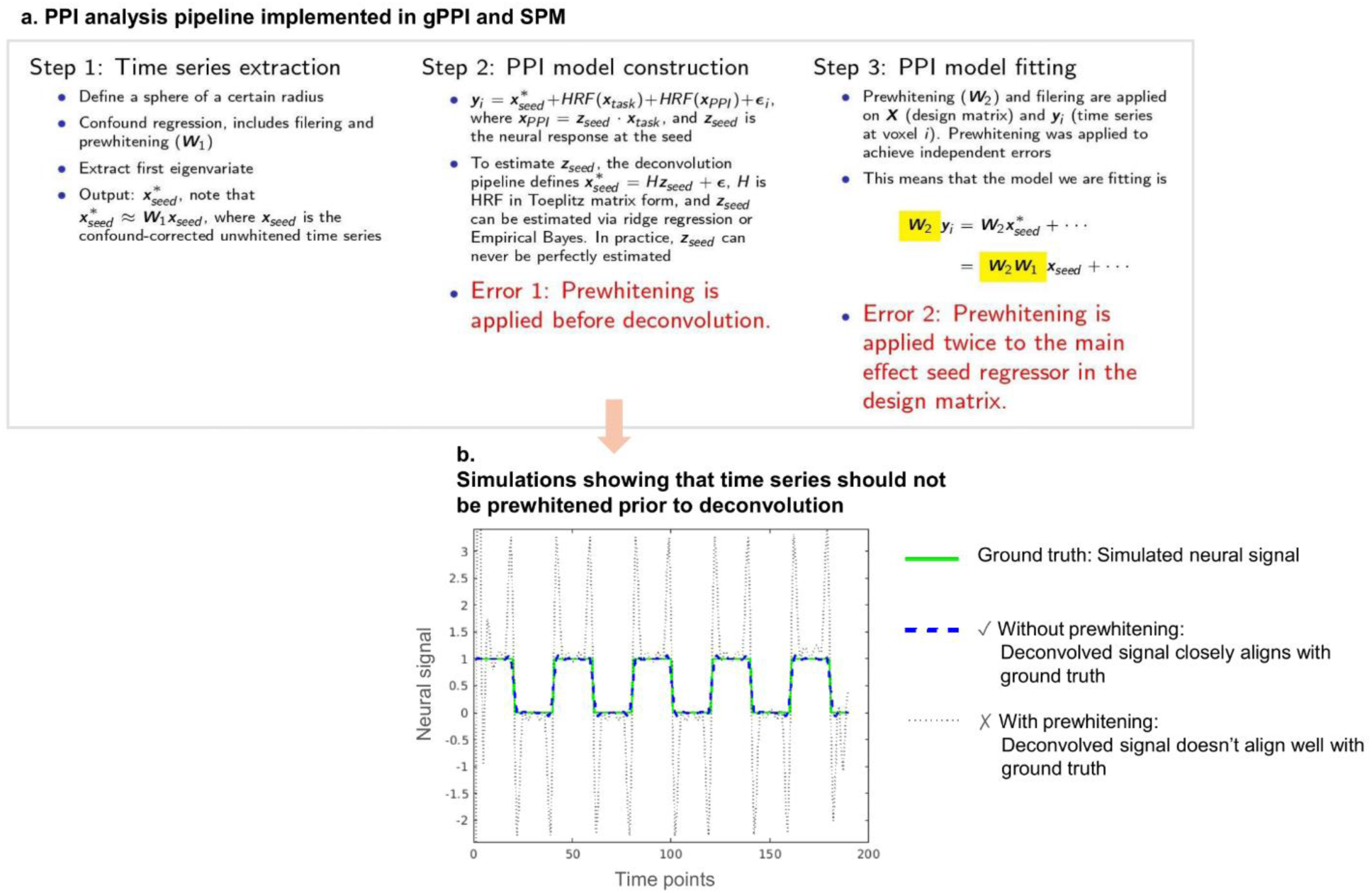
a. The PPI processing pipeline in SPM and gPPI has two issues associated with prewhitening during time series extraction. The first arises during the construction of the interaction (PPI regressor), as deconvolution is applied to a prewhitened time series. The second issue occurs during model fitting, where prewhitening is applied again to the design matrix, resulting in the main effect seed time series being prewhitened twice. b. A simulated example showing that time series should not be prewhitened prior to deconvolution, as prewhitening alters the structure of the time series. The deconvolved signal with prewhitening (grey dotted line) has been scaled and vertically shifted to better match the ground truth.

Another issue confounding PPI analyses concerns (lack of) mean-centring of the task regressor during creation of the interaction term, as well explained by Di et al. (2017). Due to the necessarily imperfect deconvolution, without mean-centring the interaction term carries unwanted correlation with the main effect of the seed. The model might incorrectly attribute this correlation to the interaction term, resulting in a spuriously inferred interaction effect (Di et al., 2017). To resolve this issue, one has to mean-centre the task regressor when calculating the interaction term (Di et al., 2017; Masharipov et al., 2024). For simplicity, we refer to this as ‘mean-centring’ in this paper.

However, a widely used software package for PPI analysis, the gPPI toolbox, with its most recent update in 2014 (version 13.1), has not incorporated the mean-centring approach advocated by Di et al. (2017). Previously, we modified the gPPI toolbox to apply PPI analysis with mean-centring to estimate task-dependent connectivity in the reading network (He et al., 2025). Here, we use this same dataset to extend the work of Di et al. (2017) to investigate how prewhitening and non-mean-centring affect group comparisons and behavioural correlations in PPI analyses. In addition, we systematically reviewed PPI literature published since 2017 to determine whether mean-centring has become a new standard in the field. Finally, in the discussion, we provide recommendations for the implementation, and reporting of PPI studies.

## 2 Methods

### 2.1 Participants

Participant characteristics have been described in He et al. (2025). Briefly, the study included 94 adult participants with seizure disorders (median age 33, interquartile range = 19; 47 males) and 107 healthy controls (median age 43, interquartile range = 22; 38 males) from the Australian Epilepsy Project (AEP). The study was approved by the Austin Health Human Research Ethics Committee (HREC/60011/Austin-2019 and HREC/68372/Austin-2022).

### 2.2 MRI data acquisition, preprocessing, and fMRI paradigm

We collected T1-weighted as well as Multi-Band Multi-Echo (MBME) fMRI images for all participants in a 3T Siemens PrismaFit MRI scanner. Details of scanning parameters and preprocessing are discussed in our previously published paper (He et al., 2025). All participants completed a block design language fMRI task. In the task-active phase, participants decide whether visually presented pairs of pseudowords rhyme or not. In the baseline phase, participants decide whether pairs of patterns of forward and backward slashes are identical. Each stimulus pair was displayed for 4.5 seconds, and each block consisted of four stimulus pairs. The experiment lasted 180 seconds (TR = 0.9 s; 200 TRs in total) and included 20 rhyming and 20 pattern matching pairs.

### 2.3 Task activation analysis

We first fitted a GLM to contrast the pseudoword rhyming blocks (coded as 1) against the pattern matching baseline (coded as 0). Confound regressors included 24 head motion parameters (Friston et al., 1996), the first eigenvariates of white matter and cerebrospinal fluid signals, and a constant term. The model also incorporated a 128-second high-pass filter and prewhitening with a FAST model, recommended for TR shorter than 1 second (Corbin et al., 2018; Olszowy et al., 2019). One sample *t*-tests were conducted on individual first level task activation estimates (n = 201). A two-tailed Family-Wise-Error cluster corrected (FWEc) threshold of p < 0.05 was applied, with an initial cluster forming threshold of p < 0.001 uncorrected. Task activation analysis was performed using the iBT software version 3.9 (Abbott et al., 2024) with SPM12 revision 7771 (Penny et al., 2006).

### 2.4 Simulations of prewhitening prior to deconvolution

To illustrate the effect of prewhitening before deconvolution, we performed a simple simulation. We generated a time series resembling a block design (green solid line in Fig. 1b) and convolved it with SPM’s canonical HRF to approximate a BOLD response. We then applied two approaches: (1) direct deconvolution of the time series and (2) prewhitening and then deconvolution. For simplicity, we estimated the whitening matrix by fitting an AR(1) model using MATLAB’s *arima* command. In the simulations, deconvolution was performed using ridge regression with a shrinkage parameter of 0.002, as this value approximates SPM’s built-in Empirical Bayes procedure (Masharipov et al., 2024).

### 2.5 Task modulated connectivity analysis

We next performed PPI analyses (Friston et al., 1997) seeding from left fusiform gyrus (FusG) to test the hypothesis that across all participants, there would be upregulation of information flow between FusG and major language nodes during pseudoword rhyming relative to pattern matching. We defined the seed region using a sphere of 6mm radius, centred on the peak activation coordinate in left FusG from the one sample *t*-test modelling the task activation (green spheres in Fig. 2 and Fig. 3). We then extracted the first eigenvariate of the time series across all voxels within the spherical region of interest (ROI) for each participant using SPM’s time series extraction function. This included applying a high-pass filter of 128 s, confound-correction, and prewhitening. All analyses were performed using SPM12 (revision 7771) and the gPPI toolbox (v13.1) with modified code (details below).

**Figure 2.**
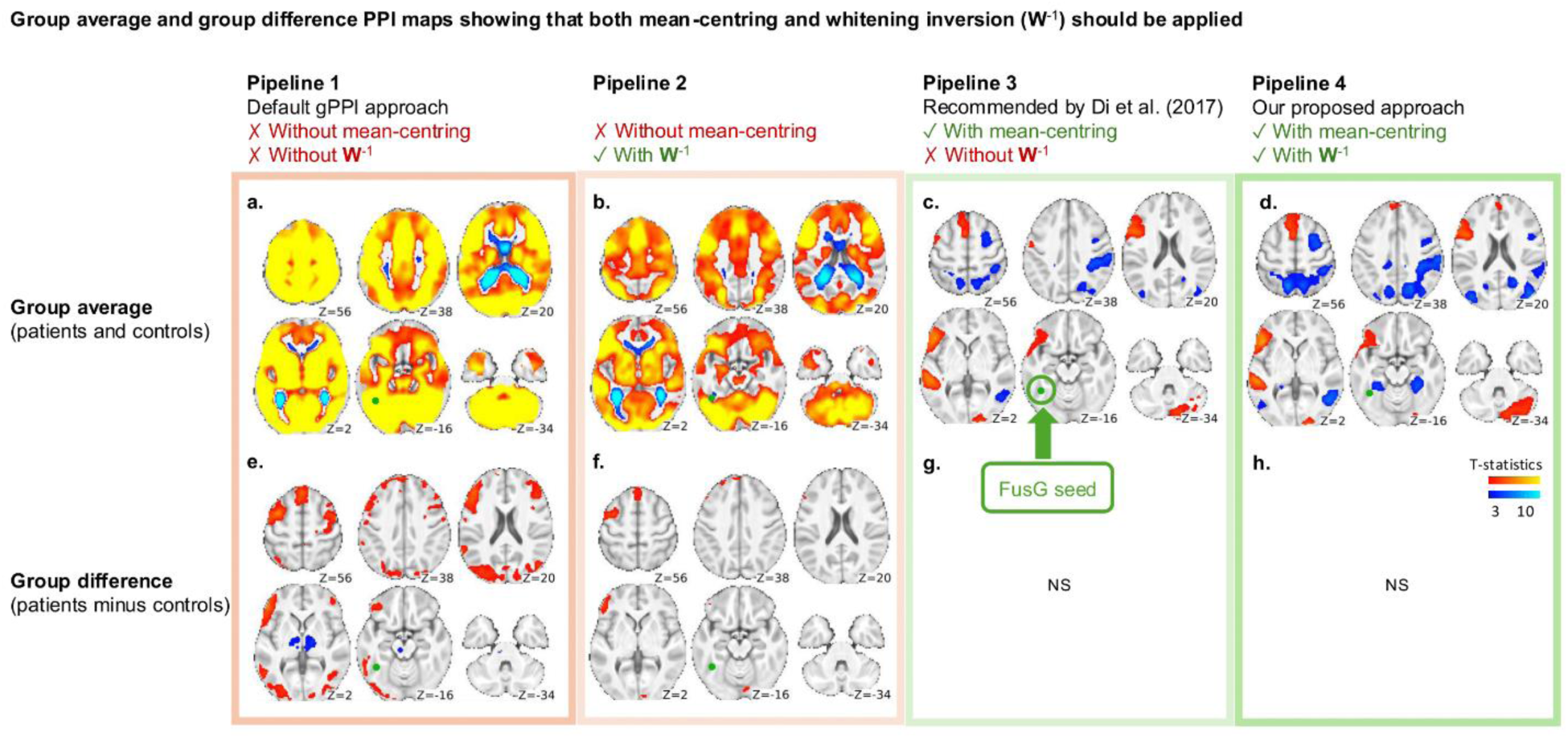
Effects of mean-centring and whitening inversion on group PPI results. Different pipelines are shown as columns. Group average and group differences in interaction effects are shown in the top and bottom rows, respectively. FusG seed location is shown in green. Left hemisphere on the left side. FWEc p < 0.05, two-sided. NS: not significant. a. Without mean-centring and without whitening inversion (Pipeline 1), leads to unreasonably widespread and implausible interaction effects, including artefactual interaction effects at the seed location. b. Whitening inversion (Pipeline 2) removes interaction effects at the seed location, but implausible widespread effects are still present. c. Mean-centring (Pipeline 3) eliminates widespread interaction effects, leaving positive effects in left language areas and negative effects in right hemisphere attentional areas. d. Combining mean-centring with whitening inversion (Pipeline 4) improves detection of interaction effects relative to mean-centring alone. Bottom row: with correct modelling (g) there are no significant differences observed between our particular groups. If only inspecting between group maps (e,f), model misspecification would not necessarily be as apparent as seen in a,b.

**Figure 3.**
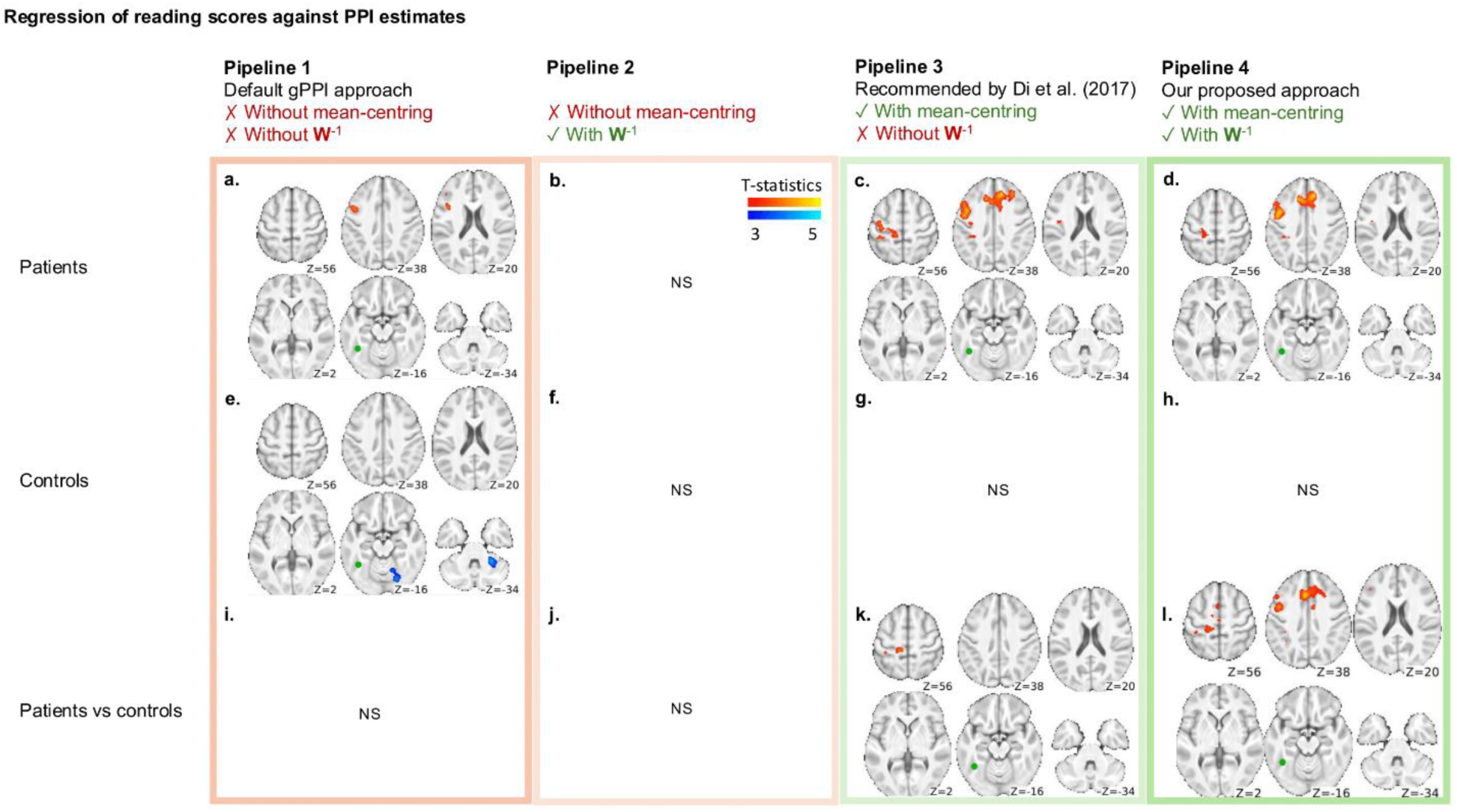
Regression of reading scores against PPI estimates. Different pipelines are shown as columns. The reference approach is to apply both mean-centring and whitening inversion (Pipeline 4). Results from Pipeline 3 (mean-centring only) were mostly consistent with Pipeline 4. Without mean-centring (Pipelines 1 and 2), results were largely inconsistent compared to Pipeline 4. However, since neither the widespread effects nor effects at the seed location were present (see Fig. 2 and Results 3.2), it may not be obvious that these results were incorrect unless a proper analysis incorporating mean-centring is performed. FusG seed location is shown in green. Left hemisphere on the left side. FWEc p < 0.05, two-sided.

#### 2.5.1 Inversion of the whitening step and mean-centring

Both whitening inversion and mean-centring are not available within the current release version of the gPPI package (v13.1) at the time of writing. We therefore implemented them by modifying the relevant lines of code. For whitening inversion, we modified the *timeseries_extract* function called at line 463 of the *PPPI.m* function, changing line 397 from *xY.u = Y* to *xY.u = inv(SPM.xX.W)*Y*. For mean-centring, we modified line 705 in *PPPI.m* function, changing *PSYxn(:,j) = PSY(:,j).*xn* to *PSYxn(:,j) = (PSY(:,j)-mean(PSY(:,j))).*xn*. Note that we only need to modify the code to mean-centre the task regressor in the interaction term, as the seed regressor is mean-centred by default.

We repeated the same PPI analysis both with and without whitening inversion, and with and without mean-centring. This resulted in four analysis pipelines:

**Pipeline 1**: Without whitening inversion and without mean-centring. This is the default pipeline used in the gPPI toolbox, and in PPI analysis implemented in SPM5 r3271 – SPM12 r6556.

**Pipeline 2**: With whitening inversion and without mean-centring.

**Pipeline 3**: Without whitening inversion and with mean-centring. This is the pipeline used in our previously published study (He et al., 2025) and is the approach recommended by Di et al. (2017). It is also the approach that would likely result from manual construction of a PPI analysis (i.e. without the gPPI toolbox) in SPM12 r7771 or later, as the regressors are mean-centred by default.

**Pipeline 4**: With whitening inversion and with mean-centring. This is our recommended pipeline.

#### 2.5.2 Second level analysis

For each pipeline, one sample t-tests were performed on group averaged interaction effects in the seizure group (n = 94), controls (n = 107), and seizure and controls combined (n = 201). We also ran a two sample t-test to compare PPI maps of the seizure group and controls. Finally, we ran a whole brain mixed effects regression model designed to investigate whether PPI parameter estimates covary with reading performance, and whether these relationships differ between the seizure group (n = 89, 2 seizure participants had missing reading scores and 3 had invalid scores) and controls (n = 94, 13 controls had invalid reading scores). Age, sex, and a constant term were included in the model as regressors of no interest. We investigated both positive and negative effects, hence a two-tailed FWEc threshold of p < 0.05 was applied, with an initial cluster forming threshold of p < 0.001 uncorrected.

### 2.6 A systematic survey on PPI literature from 2018 to 2022

We investigated whether mean-centring the task regressor during the generation of the interaction term, as recommended by Di et al (2017), has become a new standard in the field. We searched the PPI literature for articles published between 1 January 2018 and 31 December 2022 on Clarivate Web of Science (https://www.webofscience.com/wos/alldb/basic-search) that have the terms “psychophysiological interaction” and “fMRI” in their title, abstract, or keywords. A total of 169 papers were identified. We excluded 5 papers: 3 due to their focus on methodology rather than the application of PPI on an actual dataset, and 2 because the authors did not apply PPI to their data. The final number of included papers is 164.

For each paper, we screened the methods and results to categorise them into one of three categories: “unlikely affected” by the non-mean-centring issue in PPI (see Results and Di et al., 2017), “likely affected” by the non-mean-centring issue, or “unable to determine”. The screening process is as follows (also see Fig. 4):

**Figure 4.**
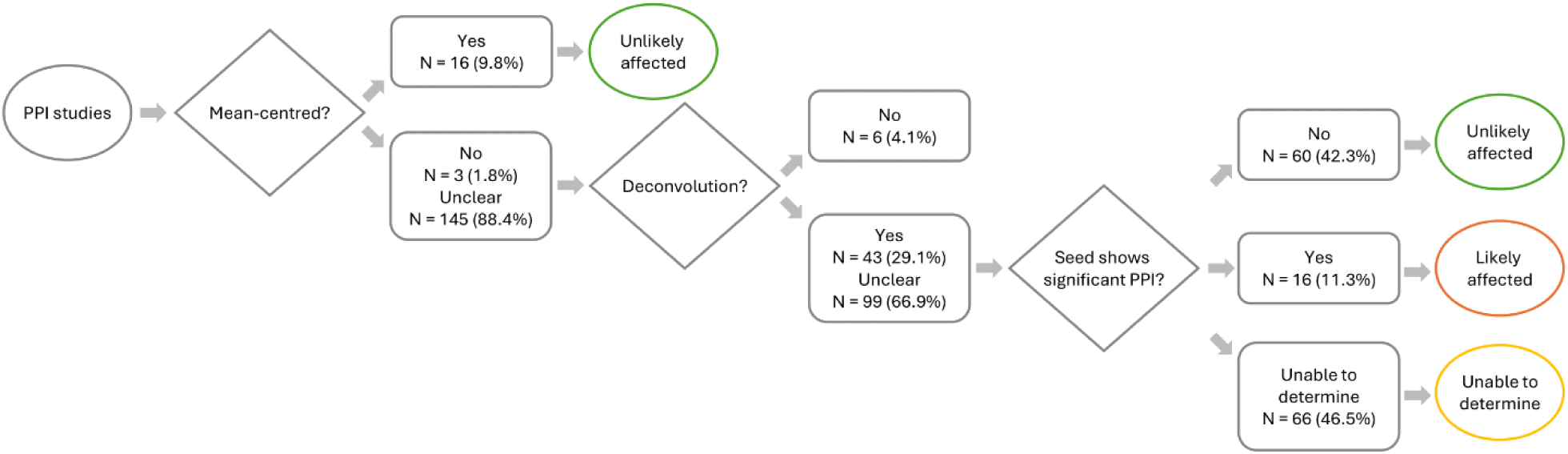
Screening flowchart. We first screened the methods section of each paper to determine whether the authors mean-centred the task regressor when calculating the interaction term. For those that did not mean centre or had unclear methods, we checked whether they applied deconvolution in their PPI analysis. For those that applied deconvolution or had unclear descriptions, we screened their results for any suspicious PPI findings. Specifically, we checked whether the reported interaction effects overlapped with the seed region itself. As discussed in Results 3.2, without mean-centring or whitening inversion, it is likely that the resulting interaction effects will peak at the seed location. We categorised each study as “unlikely affected”, “unable to determine”, or “likely affected” by the mean-centring issue. Percentages add up to 100% for each branch.

Step 1: We screened the methods to determine whether they mean-centred the task regressor when calculating the interaction term. For those that explicitly mentioned a mean-centring step, we categorised them as “unlikely affected” by the non-mean-centring issue.

Step 2: For the remaining studies, we checked whether they applied deconvolution in their PPI computations (as discussed in Di et al. (2017), mean-centring is not necessary if deconvolution is not performed.). Deconvolution is not implemented in FSL (Jenkinson et al., 2012), BrainVoyager (Goebel, 2012), and the CONN toolbox (Whitfield-Gabrieli & Nieto-Castanon, 2012). Therefore, studies using only these software were categorised as “unlikely affected”.

Step 3: For studies that did not apply mean-centring (or had unclear methods) but applied deconvolution (or had unclear methods), we screened their results for any suspicious PPI findings. Specifically, we checked whether the reported interaction effects overlapped with the seed region itself. At the seed location, the principal effect should just be that of the seed regressor. Studies that showed significant group-averaged interaction effects in the seed region were therefore categorised as “likely affected”. Studies that did not show significant group-averaged interaction effects in the seed region were categorised as “unlikely affected”. Studies that consisted of group comparisons and did not report group-averaged PPI effect maps were categorised as “unable to determine”, because some or all of the effect on the seed region might cancel out (as we show using our own data in Results 3.2). Studies that consisted of behavioural correlations and did not report group-averaged PPI effect maps were also categorised as “unable to determine”, because the effect on the seed region might not be captured in the behavioural regressor (see Results 3.2). Studies that consisted of ROI-to-ROI analysis and did not report the correlation with the seed itself were categorised as “unable to determine”. Studies that reported group-averaged PPI maps but were unclear about the location of the seed in relation to the PPI maps were categorised as “unable to determine”.

## 3 Results

### 3.1 Prewhitening adversely affects deconvolution-based estimation of the underlying neuronal signal

We first demonstrated, via simulations, that prewhitening adversely affects the deconvolution process (Fig. 1b). With whitening inversion, which is equivalent to removing prewhitening in this case, the deconvolved signal closely aligns with the simulated neural signal.

### 3.2 Group level results

#### 3.2.1 Whitening inversion removes spurious interaction effects at the seed location even without mean-centring

Without mean-centring (Pipelines 1 and 2), we observed widespread group-averaged positive interaction effects extending well beyond areas that might reasonably be hypothesised to be involved (Fig. 2a,b). The spurious effects are lessened with whitening inversion (Pipeline 2, Fig 2b), which also eliminates artefactual interaction effects at the seed location.

#### 3.2.2 Mean-centring eliminated widespread interaction effects, while whitening inversion enhanced the clusters

With mean-centring (Pipelines 3 and 4), the widespread spurious interaction effects seen in Pipelines 1 and 2 were eliminated. Significant interactions during rhyming were limited to language regions, and during pattern matching to right parietal and posterior temporal regions (Fig. 2c,d). In addition, there is no between group difference in interaction effects (Fig. 2g,h). Without mean-centring (Pipelines 1 and 2), the strong and widespread interaction effects present in each group individually (see Supplementary Fig. 1 for group specific results) largely cancel out, as do the artefactual effects at the seed location; some spurious group differences nevertheless remain (Fig. 2e,f). In other words, if only examining between group effects, the consequences of model misspecification may not be obvious.

#### 3.2.3 Impact of PPI model misspecification on behavioural regression

Taking Pipeline 4 as the reference processing approach, we found a positive relationship between reading scores and the PPI parameter estimates in the superior medial frontal region and the left middle frontal gyrus, in the seizure patient group (Fig. 3d). In contrast, no such relationship was observed in controls (Fig. 3h), and the difference was significant between patients and controls (Fig. 3l). If only mean-centring was applied without whitening inversion (Pipeline 3), the results appeared broadly comparable (Fig. 3c,g), though the group differences diminish (Fig. 3k). Without mean-centring (Pipelines 1 and 2), results are either missed (Fig. 3b), diminished (Fig. 3 a) or spurious (Fig. 3 e).

### 3.3 PPI mean-centring error remains an issue in 10% or more of published studies

We have shown that in deconvolution-based PPI models mean-centring reduces the likelihood of spurious interaction effects and whitening inversion increases power to detect true interaction effects. Given Di alerted the field to the mean-centring issue in 2017, and the risk of false positive findings from failing to mean-centre, we surveyed PPI studies published between 2018 and 2022 to investigate contemporary adoption of the mean centring approach. Among the 164 studies identified, we categorised 16 (9.8 %) as “likely affected”, 66 (40.2 %) as “unable to determine”, and 82 (50.0 %) as “unlikely affected” by this non-mean-centring issue in PPI. Thus, 10% or more of recently published studies using PPI methods are likely based upon misspecified models (Fig. 4).

## 4 Discussion

Through both simulation and the analysis of real data we have demonstrated two issues in the implementation of PPI analysis in SPM and the gPPI toolbox, one previously reported - mean-centring, and the other novel - prewhitening. The practical consequences of these two issues on group average interaction effects are highlighted in Fig. 2a-d. Comparing Fig. 2a or Fig. 2b with Fig. 2c demonstrates that mean centring is essential to avoid false-positive PPI effects, regardless of the prewhitening issue. We expect this, because mean-centring ensures there is no main effect seed component in the PPI term (Di et al., 2017). However, comparing Fig. 2a with Fig. 2b reveals prewhitening can worsen the false positive PPI effects that result from a lack of mean-centring, including emergence of false positive PPI at the seed location. Finally, comparing Fig. 2c with Fig. 2d demonstrates the optimum model (Pipeline 4; where the seed regressor is better represented in the model) confers additional power to detect true PPI effects.

We further showed that group comparisons can conceal some of these effects, and that model misspecification has deleterious consequences for detecting behavioural correlations. In a systematic survey of the recent literature we discovered that PPI model misspecification remains a non-negligible problem in contemporary work.

### 4.1 Presence of a significant interaction effect at the seed location as a ‘red flag’

The presence of a significant interaction effect at the seed location itself is a telltale sign of model misspecification. We have shown that including a whitening inversion step ensures that the main effect seed regressor in the PPI model is well-matched to the signal at the seed location itself, preventing the emergence of spurious interaction effects at the seed location (Fig. 2b). We note that an alternative approach is to skip prewhitening in time series extraction, as the regression coefficients in confound regression should remain asymptotically unbiased despite the presence of autocorrelation (Gauss-Markov Theorem; see also Arbabshirani et al., 2014; Penny et al., 2006; and Olszowy et al., 2019). However, skipping this step would require significant code changes, as SPM’s confound regression relies on prewhitened first level beta maps. Alternatively, one may achieve this by additionally performing a first level analysis without specifying an autocorrelation model and then extract time series from its output.

### 4.2 Consequences of not mean-centring

If mean-centring is not performed, the interaction term carries a seed component that does not match perfectly with the main effect seed regressor due to the deconvolution algorithm, whereas mean-centring avoids this component (Di et al., 2017). To understand the potential consequences of mean-centring or not, we first compared the patterns of results that follow from these two approaches when applied to our own data. Using a deconvolution approach with proper mean-centring, we observed task dependent connectivity from left FusG to language and visuospatial attentional regions, in both seizure and control participants (Fig. 2; details in He et al., 2025). Without mean-centring, none of the PPI maps are valid due to the extra unwanted variance in the interaction term that highly correlates with the main effect of the seed (Fig. 2 and 3). The consequences of the misspecification are widespread and obvious when we look at group-averaged interaction effects, but less obvious in the group comparison map and the second level regression maps (first two columns Fig. 2 and 3) - it becomes evident that these maps cannot be trusted when compared to the correctly mean-centred results (last two columns Fig. 2 and 3).

### 4.3 Survey of contemporary practice

In the systematic survey of recently published PPI literature, we noticed many studies fall into the “unable to determine” category due to insufficient details provided in the paper. For example, many papers neither indicated whether mean-centring was applied nor specified how the task regressor was coded. Certain versions of SPM, and the gPPI toolbox, do not apply mean-centring in PPI models, and it is impossible to extract this information if the software version number is not included in the paper. As discussed in Results 3.2.1, without mean-centring and without whitening inversion, the seed will show a strong interaction effect. Therefore, for studies with unclear methods regarding whether mean-centring and/or deconvolution were applied, we needed to screen their results to check whether the reported interaction effects overlap with the seed. During this process, often we found it unclear where the seed is located on the PPI maps. We also note that comparing groups or performing second level regressions without considering the PPI results at the single group level can make it challenging to detect abnormalities. In addition, some studies adopted a ROI-to-ROI approach to increase power, as the effect size of the interaction term may be much smaller than that of the main effects (O’Reilly et al., 2012). In this case, if the authors do not check the interaction with the seed region itself, the results can be difficult to assess.

As a side note, we noticed that the modal sample size used in the 164 PPI papers reviewed in this study was 23 (IQR = 19.75). Based on our own analysis, the interaction term has an effect size of approximately 0.4 (calculated by dividing the mean beta estimates across each cluster by the standard deviation). To achieve power of 0.8, at least 52 participants are required (Faul et al., 2007), which is consistent with a simulation-based study by Masharipov et al. (2024). Therefore, most of the included studies may be underpowered. Without proper mean-centring, the significant results could be borrowing from the main effects.

### 4.4 Recommendations for implementing and reporting PPI analysis

We propose several recommendations for conducting PPI analyses. Firstly, it is important for users to understand how an analysis has been implemented in the software being used. In the context of PPI, one should be aware of how a time series is extracted, as well as whether mean-centring has been applied by default and whether a deconvolution step is involved. After implementing a PPI analysis, a quick sanity check is to screen the interaction effect in the seed region. A misspecified PPI model is likely to show a significant interaction effect in the seed, as discussed in Results 3.2.1 and Di et al. (2017). For group comparisons, one should check the PPI maps at a single group level. For ROI-to-ROI analysis, one should check the interaction effects with the seed ROI itself. It is also important to note that non-mean-centred PPI results may consist of a combination of correct and spurious interaction effects. It would therefore be challenging to disentangle these effects without conducting a proper PPI analysis with mean-centring.

When reporting a PPI analysis, authors should detail whether mean-centring and deconvolution were applied, as well as the software version used to implement the analysis (Nichols et al., 2016). In the results section, authors should report their PPI maps with reference to location of the seed (e.g. by showing slices that intersect with the seed, and with the seed indicated). In studies with multiple group comparisons or second level regressions, it may be useful to also include single group PPI maps even if only as supplementary information. At the very least, authors should review their own single group maps as a data quality control step, to check for spurious interaction effects that intersect with the seed location – a sure sign of model misspecification. In studies with ROI-to-ROI analysis, authors should also report the interaction effect with the seed ROI.

### 4.5 Conclusion

In conclusion, we identified two issues associated with conventional PPI approaches that use deconvolution: one with prewhitening and one with mean-centring. We demonstrated that these errors are non-negligible through both simulations and applications to empirical data. Notably, such model misspecification can be easily overlooked if one examines only between group interaction effects. Additionally, a systematic survey of recent literature revealed that model misspecification and insufficient methods reporting remain significant issues in the field. Our findings show that PPI is a valuable technique for interrogating brain networks in vivo, but highlight the importance of proper model specification to ensure appropriate inference.

## 5 Data and code availability

Any requests for access to the data used in this project should be directed to the Australian Epilepsy Project (a formal data access request can be lodged at https://www.epilepsyproject.org.au/research/access-to-aep-data). The analyses were conducted using publicly available software, and no custom code was developed other than the two modifications described in the manuscript.

## 6 Author contributions

V. He: Conceptualisation, Methodology, Formal Analysis, Writing - Original Draft, Writing - Review & Editing, Visualisation. B. Tahayori: Resources, Data Curation, Writing - Review & Editing. D.N. Vaughan: Conceptualisation, Methodology, Investigation, Resources, Writing - Review & Editing, Project Administration, Funding Acquisition. H.R. Pardoe: Investigation, Resources, Writing - Review & Editing, Project Administration. G.D. Jackson: Conceptualisation, Methodology, Investigation, Resources, Writing - Review & Editing, Supervision, Project Administration, Funding Acquisition. C. Tailby: Conceptualisation, Methodology, Investigation, Resources, Data Curation, Writing - Original Draft, Writing - Review & Editing, Visualisation, Supervision, Project Administration, Funding Acquisition. D.F. Abbott: Conceptualisation, Methodology, Software, Investigation, Resources, Data Curation, Writing - Original Draft, Writing - Review & Editing, Visualisation, Supervision, Project Administration, Funding Acquisition. The full list of AEP investigators is presented in Supplementary Table 1.

## 8 Acknowledgements

The authors thank Karl J. Friston and Peter Zeidman from the Functional Imaging Laboratory at UCL for their endorsement of this work. The Australian Epilepsy Project received funding from the Australian Government under the Medical Research Future Fund (Frontier Health and Medical Research Program - Grant Numbers MRFF75908 and RFRHPSI000008) and the Victoria State Government (Victorian-led Frontier Health and Medical Research Program). The Florey Institute of Neuroscience and Mental Health acknowledges the strong support from the Victorian Government and in particular the funding from the Operational Infrastructure Support Grant. The authors acknowledge the facilities and scientific and technical assistance of the National Imaging Facility, a National Collaborative Research Infrastructure Strategy (NCRIS) capability. This research was supported by The University of Melbourne’s Research Computing Services and the Petascale Campus Initiative. VH acknowledges the financial support received from the University of Melbourne through the Melbourne Research Scholarship. DFA and BT acknowledge fellowship funding from the National Imaging Facility.

## References

Abbott, D. F., Capon, A., Tahayori, B., Bryant, M., Vaughan, D. N., Jackson, G. D., & for the Australian Epilepsy Project Investigators. (2024, June). The Integrated Brain Analysis Toolbox for SPM (iBT). OHBM 2024 – 30th Annual Meeting of the Organization for Human Brain Mapping, Seoul, Korea. https://archive.aievolution.com/2024/hbm2401/Abstracts/viewAbs?abs=2320

Arbabshirani, M. R., Damaraju, E., Phlypo, R., Plis, S., Allen, E., Ma, S., Mathalon, D., Preda, A., Vaidya, J. G., Adali, T., & Calhoun, V. D. (2014). Impact of autocorrelation on functional connectivity. NeuroImage, 102, 294–308. 10.1016/j.neuroimage.2014.07.045

Cole, M. W., Reynolds, J. R., Power, J. D., Repovs, G., Anticevic, A., & Braver, T. S. (2013). Multi-task connectivity reveals flexible hubs for adaptive task control. Nature Neuroscience, 16(9), 1348–1355. 10.1038/nn.3470

Corbin, N., Todd, N., Friston, K. J., & Callaghan, M. F. (2018). Accurate modeling of temporal correlations in rapidly sampled fMRI time series. Human Brain Mapping, 39(10), 3884–3897. 10.1002/hbm.24218

Di, X., & Biswal, B. B. (2017). Psychophysiological Interactions in a Visual Checkerboard Task: Reproducibility, Reliability, and the Effects of Deconvolution. Frontiers in Neuroscience, 11, 573. 10.3389/fnins.2017.00573

Di, X., Reynolds, R. C., & Biswal, B. B. (2017). Imperfect (de)convolution may introduce spurious psychophysiological interactions and how to avoid it. Human Brain Mapping, 38(4), 1723–1740. 10.1002/hbm.23413

Faul, F., Erdfelder, E., Lang, A.-G., & Buchner, A. (2007). G*Power 3: A flexible statistical power analysis program for the social, behavioral, and biomedical sciences. Behavior Research Methods, 39(2), 175–191. 10.3758/BF03193146

Friston, K. J., Buechel, C., Fink, G. R., Morris, J., Rolls, E., & Dolan, R. J. (1997). Psychophysiological and modulatory interactions in neuroimaging. NeuroImage, 6(3), 218–229. 10.1006/nimg.1997.0291

Friston, K. J., Williams, S., Howard, R., Frackowiak, R. S., & Turner, R. (1996). Movement-related effects in fMRI time-series. Magnetic Resonance in Medicine, 35(3), 346–355. 10.1002/mrm.1910350312

Gitelman, D. R., Penny, W. D., Ashburner, J., & Friston, K. J. (2003). Modeling regional and psychophysiologic interactions in fMRI: The importance of hemodynamic deconvolution. NeuroImage, 19(1), 200–207. 10.1016/s1053-8119(03)00058-2

Goebel, R. (2012). BrainVoyager—Past, present, future. NeuroImage, 62(2), 748–756. 10.1016/j.neuroimage.2012.01.083

He, et al. (2025). Task demands shape network interactions during reading and visual form processing. Imaging Neuroscience, in press.

Jenkinson, M., Beckmann, C. F., Behrens, T. E. J., Woolrich, M. W., & Smith, S. M. (2012). FSL. NeuroImage, 62(2), 782–790. 10.1016/j.neuroimage.2011.09.015

Masharipov, R., Knyazeva, I., Korotkov, A., Cherednichenko, D., & Kireev, M. (2024). Comparison of whole-brain task-modulated functional connectivity methods for fMRI task connectomics. Communications Biology, 7(1), 1–21. 10.1038/s42003-024-07088-3

McLaren, D. G., Ries, M. L., Xu, G., & Johnson, S. C. (2012). A generalized form of context-dependent psychophysiological interactions (gPPI): A comparison to standard approaches. NeuroImage, 61(4), 1277–1286. 10.1016/j.neuroimage.2012.03.068

Nichols, T. E., Das, S., Eickhoff, S. B., Evans, A. C., Glatard, T., Hanke, M., Kriegeskorte, N., Milham, M. P., Poldrack, R. A., Poline, J.-B., Proal, E., Thirion, B., Essen, D. C. V., White, T., & Yeo, B. T. T. (2016). *Best Practices in Data Analysis and Sharing in Neuroimaging using MRI* (p. 054262). bioRxiv. 10.1101/054262

Olszowy, W., Aston, J., Rua, C., & Williams, G. B. (2019). Accurate autocorrelation modeling substantially improves fMRI reliability. Nature Communications, 10(1), Article 1. 10.1038/s41467-019-09230-w

O’Reilly, J. X., Woolrich, M. W., Behrens, T. E. J., Smith, S. M., & Johansen-Berg, H. (2012). Tools of the trade: Psychophysiological interactions and functional connectivity. Social Cognitive and Affective Neuroscience, 7(5), 604–609. 10.1093/scan/nss055

Park, H. J., & Friston, K. J. (2013). Structural and functional brain networks: From connections to cognition. *Science (New York*, N.Y.), 342(6158), 1238411. 10.1126/science.1238411

Penny, W. D., Friston, K. J., Ashburner, J. T., Kiebel, S. J., Nichols, T. E., & Nichols, T. E. (2006). Statistical Parametric Mapping: The Analysis of Functional Brain Images. Elsevier Science & Technology. http://ebookcentral.proquest.com/lib/unimelb/detail.action?docID=282095

Whitfield-Gabrieli, S., & Nieto-Castanon, A. (2012). Conn: A functional connectivity toolbox for correlated and anticorrelated brain networks. Brain Connectivity, 2(3), 125–141. 10.1089/brain.2012.0073

