## Supplementary Information for "Common pitfalls during model specification in psychophysiological interaction analysis"

**List of Contents**

**Supplementary Figure 1**

**Supplementary Table 1**

**Supplementary Figure 1**

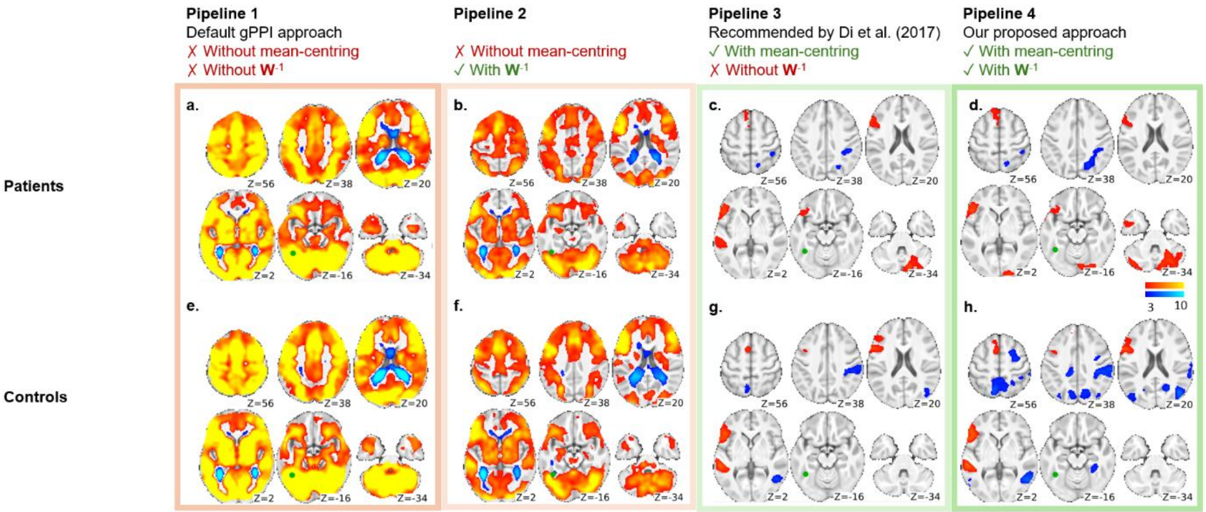


Fig. 1. Effects of mean-centring and whitening inversion on group specific PPI results. Different pipelines are shown as columns. Patient group and control group interaction effects are shown in the top and bottom rows, respectively. FusG seed location is shown in green. Left hemisphere on the left side. FWEc $p < 0.05$, two-sided. Direct comparisons of patients versus controls are shown in the manuscript Figure 2; no significant between group differences are observed when using Pipelines 3 or 4.

Table 1. AEP investigator list with Contributor Roles Taxonomy (CRediT) author statement relevant for this manuscript.

| Australian Epilepsy Project Investigators | | | |
|---|---|---|---|
| **Name & ORCID** | **Primary Location** | **Role** | **CRediT** |
| **Graeme D. Jackson, MD 0000-0002-7917-5326** | The Florey Institute of Neuroscience and Mental Health | Chief Investigator | Conceptualisation, Methodology, Investigation, Resources, Writing - Review & Editing, Supervision, Project Administration, Funding Acquisition |
| **David F. Abbott, PhD 0000-0002-7259-8238** | The Florey Institute of Neuroscience and Mental Health | Informatics Lead | Conceptualisation, Methodology, Software, Investigation, Resources, Data Curation, Writing - Original Draft, Writing - Review & Editing, Visualisation, Supervision, Project Administration, Funding Acquisition |
| **Zanfina Ademi, PhD** | Monash University | Health Economics Lead | Conceptualisation, Funding acquisition |

| | | | |
|---|---|---|---|
| **0000-0002-0625-3522** | | | |
| **Subhaga Amarasekara** | The Florey Institute of Neuroscience and Mental Health | Product Lead | Resources, Project Administration |
| **Amanda Anderson** | The Florey Institute of Neuroscience and Mental Health | Lived Experience Ambassador and Participant Lead | Investigation, Resources, Funding acquisition |
| **Rachel Hughes** | The Florey Institute of Neuroscience and Mental Health | Clinical Research Coordinator | Investigation, Resources |
| **Donna Hutchison** | The Florey Institute of Neuroscience and Mental Health | Executive Lead | Project administration |
| **Patrick Kwan, MD 0000-0001-7310-276X** | Monash University | Outcomes Lead | Conceptualisation, Resources, Funding acquisition |
| **Paul Lightfoot** | The Florey Institute of Neuroscience and Mental Health | Operations Lead | Investigation, Project administration |
| **Saul Mullen, MD, PhD 0000-0003-1224-4101** | The University of Melbourne | Protocol Development Lead (2019-2021) | Conceptualisation, Methodology, Funding acquisition |
| **Karen L. Oliver, PhD** | The University of Melbourne | Genetics Lead | Conceptualisation, Funding acquisition |

| | | | |
|---|---|---|---|
| **0000-0001-5188-6153** | | | |
| **Heath R. Pardoe, PhD 0000-0002-0123-2167** | The Florey Institute of Neuroscience and Mental Health | Science Operations Lead | Investigation, Resources, Writing - Review & Editing, Project Administration |
| **Mangor Pedersen, PhD 0000-0002-9199-1916** | Auckland University of Technology | Artificial Intelligence Lead | Conceptualisation, Methodology, Funding acquisition |
| **Chris Tailby, PhD 0000-0002-1320-5924** | The Florey Institute of Neuroscience and Mental Health | Neuropsychology Lead | Conceptualisation, Methodology, Investigation, Resources, Data Curation, Writing - Original Draft, Writing - Review & Editing, Visualisation, Supervision, Project Administration, Funding Acquisition |
| **David N. Vaughan, MD, PhD 0000-0002-6225-7739** | The Florey Institute of Neuroscience and Mental Health | Imaging Lead | Conceptualisation, Methodology, Investigation, Resources, Writing - |

| | | | |
|---|---|---|---|
| | | | Review & Editing, Project Administration, Funding Acquisition |
| **Anton De Weger** **0009-0006-7478-361X** | The Florey Institute of Neuroscience and Mental Health | Digital and Technology Lead | Software, Resources, Data Curation |
